# Prediction of high-risk liver cancer patients from their mutation profile: Benchmarking of mutation calling techniques

**DOI:** 10.1101/2021.12.17.473127

**Authors:** Sumeet Patiyal, Anjali Dhall, Gajendra P. S. Raghava

## Abstract

Identification of somatic mutations with high precision is one of the major challenges in prediction of high-risk liver-cancer patients. In the past, number of mutation calling techniques have been developed that include MuTect2, MuSE, Varscan2, and SomaticSniper. In this study, an attempt has been made to benchmark potential of these techniques in predicting prognostic biomarkers for liver cancer. Initially, we extracted somatic mutations in liver-cancer patients using VCF and MAF files from the cancer genome atlas. In terms of size, the MAF files are 42 times smaller than VCF files and containing only high-quality somatic mutations. Further, machine learning based models have been developed for predicting high-risk cancer patients using mutations obtain from different techniques. The performance of different techniques and data files have been compared based on their potential to discriminate high and low risk liver-cancer patients. Finally, univariate survival analysis revealed the prognostic role of highly mutated genes. Based on correlation analysis, we selected 80 genes negatively associated with the overall survival of the liver cancer patients. Single-gene based analysis showed that MuTect2 technique based MAF file has achieved maximum HR_LAMC3_ 9.25 with p-value 1.78E-06. Further, we developed various prediction models using selected genes for each technique, and the results indicate that MuTect2 technique based VCF files outperform all other methods with maximum AUROC of 0.72 and HR 4.50 (p-value 3.83E-15). Eventually, VCF file generated using MuTect2 technique performs better among other mutation calling techniques to explore the prognostic potential of mutations in liver cancer. We hope that our findings will provide a useful and comprehensive comparison of various mutation calling techniques for the prognostic analysis of cancer patients.

## Introduction

According to the world health organization, cancer is a life-threatening disease and the first leading cause of death worldwide in 2019. Global cancer statistics estimate that in 2020, 19.3 million new cases and 10 million deaths have been occurred due to cancer [1]. Cancer is extremely heterogeneous; therefore, the same treatment strategy is not effective for individuals with similar types of cancer. Till now, there is no universal treatment available for all types of malignancies. Currently, several targeted therapies are available for cancer treatment, which majorly focus on the detection of mutations at the genetic level [2]. In the last few years, several therapies have been designed based on the mutated genes for the cancer treatment. For instance, B-Raf Proto-Oncogene, Serine/Threonine Kinase (BRAF) inhibitors (Sorafenib) is identified to treat melanoma patients with V600E mutation in the BRAF gene [3, 4]. However, drugs like afatinib and erlotinib are used to target the mutation in the EGFR in non-small-cell lung cancer [5, 6]. Moreover, BRCA1/BRCA2 gene mutations in ovarian cancer patients have been treated by poly (ADP-ribose) polymerase (PARP) inhibitor, i.e., olaparib [7]. Of note, research on the mutations associated with the genes in cancer patients is essential for identifying the correct mechanism of the disease. Due to the advancements in next-generation sequencing, such as whole-genome, whole-exome, and mutation calling techniques, the detection of more than 98% mutations associated with the disease using sequencing data is possible [8, 9]. The easy availability and low cost of next-generation sequencing techniques enable researchers to perform experiments on large cohorts of cancer patients [10].

The genetic variants are mainly categorised into single nucleotide variant (SNV), insertion/deletion (indel), and structural variants (SV, which incorporates copy number alterations, duplications, and translocations). In recent years, a huge number of somatic mutation calling algorithms (for example, Mutect2, Varscan2, SomaticSniper, MuSE, Strelka2, etc.) have been developed to identify mutations at the genetic level using sequencing data [11–17]. Mutect2 calls somatic mutation such as single nucleotide alterations and indels using the local assembly of haplotypes. SomaticSniper pipeline detects somatic SNVs using Bayesian algorithm to compare the genotype likelihoods in the tumor and normal samples. However, Varscan2 mutation calling algorithm uses exomes, whole-genome sequencing data to capture germline variants, somatic mutations and copy number variants in tumor-normal data. Moreover, MuSE is a Markov Substitution model for Evolution, to identify novel mutations in the large-scale tumor sequencing data.

Liver cancer is one of the deadliest disease which is the seventh most common cancer among the 36 cancers reported by Global Cancer Statistics 2020 [1]. Ample treatment methods were developed in the past, but still the survival rate of liver cancer patients is very low, leading to high-mortality rate [18]. Being the most comprehensive resource for the cancer related research, TCGA provides two types of file formats for mutation data such as Variant Call Format (VCF) and Mutation Annotation Format (MAF). VCF files are the raw mutation files that store and report the genomic sequence variations that directly came out of the various automated variant calling pipelines. On the other hand, MAF files are the processed version of the VCF files, which are curated by removing the false positives or by recovering the known calls that the automated pipelines may have missed. VCF files report mutations irrespective of their importance, but MAF files describe only the most affected ones by removing the low-quality mutations. In GDC portal, both type of files are available generated using the four major mutation calling techniques named as MuTect2, MuSE, Varscan2, and SomaticSniper. Despite number of techniques are available, it is difficult to understand which method and file is better to explore the role of mutations in cancer.

In the current study, we have systematically evaluated the four mutation calling tools which are widely used in TCGA, to identify highly mutated genes associated with high-risk liver cancer patients. For this, we have collected VCF and MAF files of 418 liver cancer patients for all the mutation calling techniques. The gene-based annotations were identified using highly accurate and widely used methods ANNOVAR [19] and Maftools [20]. Correlation and survival analysis is performed to identify mutated genes that can impact the survival of liver cancer patients. Finally, several prediction algorithms have been developed for the top genes. The inferences of our study can give a valuable reference and guidance to the researchers to choose a reliable somatic mutation algorithm to determine the mutation-associated genes having a significant impact on the survival of the cancer patients.

## Material and Methods

### Dataset Collection

We obtained liver cancer (TCGA-LICH and TCGA-CHOL) mutation data from Genome Data Commons (GDC) data portal. Precisely, we collected the controlled access VCF of liver cancer patients under the approval of dbGap (Project No. 17674) according to the GDC protocols [21]. In addition to that, we have also downloaded the MAF files of TCGA liver cancer patients. In TCGA, four different techniques are used for mutation calling, i.e., MuSE, Mutect2, Varscan2, and SomaticSniper. In this study, we have utilized VCF and MAF files of 418 liver cancer samples generated from four different mutation calling methods. Moreover, the clinical data like age, gender, tumor stage, overall survival (OS) time, and vital status were collected using TCGA assembler 2 [22].

### Mutation Annotations

We used the ANNOVAR software package (https://annovar.openbioinformatics.org/en/latest/) for functional annotations of genetic variant mutations. First, we convert VCF files into ANNOVAR genetic variants file; using “convert2annovar.pl” script; the processed file contains five major columns such as chromosome number, start position, end position, reference nucleotide, and altered nucleotides. It provides three major type of annotations (i.e., gene-based, region-based, and filter-based annotations). In this work, we used gene-based annotations, in which we obtained mutations/gene/samples. In this way, we get per-gene mutations for each sample for the four different mutation calling techniques. After that, we count number of mutations per gene for each liver cancer patient with the help of in-house python script (gene_to_matrix.py). Similarly, for MAF files we counted the number of mutations/gene/samples. Finally, we generated matrices for each mutation calling technique from VCF and MAF files, in which number of mutations per gene per sample were reported.

### Correlation Analysis

To understand the impact of number of genetic mutations on overall survival (OS) of liver cancer patients, we have implemented correlation test. After that, we removed the genes with the non-significant p-value i.e., >0.05, and ranked the remaining genes on the bases of correlation coefficients. We choose top-10 negatively correlated genes from each technique for VCF and MAF files for further analysis.

### Survival Analysis

In this study, we have performed survival analysis by the ‘survival’ package in R (V.3.5.1) using cox proportional hazard (Cox PH) model. We perform univariate survival, in order to understand the impact of per gene mutations on the survival of liver cancer patients. The log-rank test was used to estimate the significant survival distributions between high-risk and low-risk groups in terms of the p-value. Kaplan-Meier (KM) survival curves were used for the graphical representation of high-risk and low-risk groups [23].

#### Machine learning Techniques

##### Classification Models

In this study, we have implemented various machine learning techniques for the classification of high-risk and low-risk samples based on the number of mutations in the chosen genes. Classification algorithms includes Decision tree (DT), Support Vector Classifier (SVC), Random Forest (RF), XGBoost (XGB), Gaussian Naive Bayes (GNB), Logistic Regression (LR), k-nearest neighbors (KNNs) and ExtraTree (ET) using Scikit learn [24].

##### Regression Models

Further, we implemented several regressors to develop regression models for overall survival time prediction in liver cancer patients. These techniques were developed using python-library scikit-learn and includes Random Forest (RF), Ridge, Lasso, Decision Tree (DT), Elastic Net (ENR), Logistic Regression (LR), and Support Vector Regression (SVR)[24].

#### Performance Evaluation

##### Cross-Validation Technique

To avoid over-optimization in the machine learning models, we have used standard five-fold cross-validation technique [25, 26]. In case of classification, the complete dataset was divided into 80:20 ratio, the five-fold cross-validation was performed on the 80% training dataset. In this method, the training dataset split-up into five equal sets. However, four sets used for training and remaining set used for the testing purpose. The similar task was repeated for at least five times, so that every set can be used in training and testing. Finally, the performance or outcome computed by taking the mean of all five sets. The similar process was repeated for the cross validation of regression models. In this the complete dataset was used for the five-fold cross validation.

##### Performance Measure Parameters

To evaluate the performance of classification models, we have used standard parameters. We have calculated threshold-dependent such as sensitivity (Sens), specificity (Spec), accuracy (Acc), F1-score, and MCC, and independent parameters like Area Under the Receiver Operating Characteristic (AUROC). These parameters were calculated using the following equations (1–5).

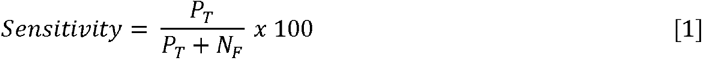

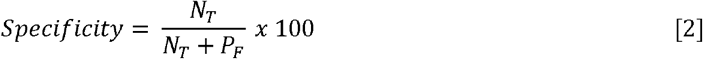

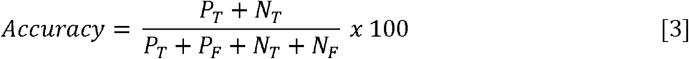

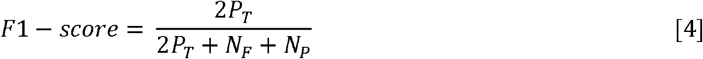

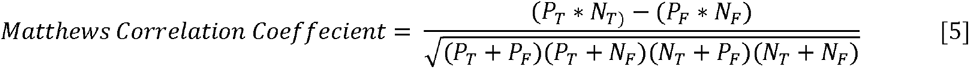

**P_T_=True Positive, P_F_=False Positive, N_T_=True Negative, N_F_=False Negative**

Similarly, to evaluate the regression models, we have used parameters such as mean absolute error (MAE), root mean-square error (RMSE), correlation coefficient (R), and p-value, to evaluate the performance of regression models as previously used in different studies [27–29].

## Results

In this study, we have used 418 TCGA liver cancer patients somatic mutation data (VCF files and MAF files) and OS data. The mutation data were taken from four different mutation calling techniques i.e., MuSE, Mutect2, Varscan2 and SomaticSniper. ANNOVAR software and in-house scripts were used to extract the number of mutations/gene/samples from the VCF and MAF files. The total number of genes and mutations extracted from different techniques is shown in Table 1. Where, in VCF files Mutect2 and SomaticSniper reported the highest number of genes and mutation counts i.e., more than 25000 genes and 5 million mutations. On the other hand, in MAF files the reported number of genes and mutations is comparatively less for each technique.

**Table 1:** Total number of genes and mutations for each gene extracted from VCF and MAF files using different mutation calling technique.

| File Type | Technique | Number of Genes | Number of Mutations |
| --- | --- | --- | --- |
| VCF | MuTect2 | 25366 | 5237093 |
|  | MuSE | 19425 | 379368 |
|  | Varscan2 | 19422 | 576231 |
|  | SomaticSniper | 25785 | 5003969 |
| MAF | MuTect2 | 16474 | 59741 |
|  | MuSE | 15712 | 51184 |
|  | Varscan2 | 15950 | 54877 |
|  | SomaticSniper | 14979 | 44102 |

Further, in order to understand the distribution of genes in each technique, we developed upset plot as shown in Figure 1. For the visualization of intersecting genes set we have created UpSet plot [30]. According to the plots, in VCF file 18758 genes were common in all the four techniques, whereas 182, 5, 2, and 630 genes are uniquely reported by MuTect2, MuSE, Varscan2, and SomaticSniper technique, respectively. Similarly, in case of MAF files 14585 genes were shared by all the techniques, while 461 genes are unique in file by MuTect2 technique, 73 by MuSE, 115 by Varscan2, and 41 unique genes were reported by SomaticSniper technique.

**Figure 1:**
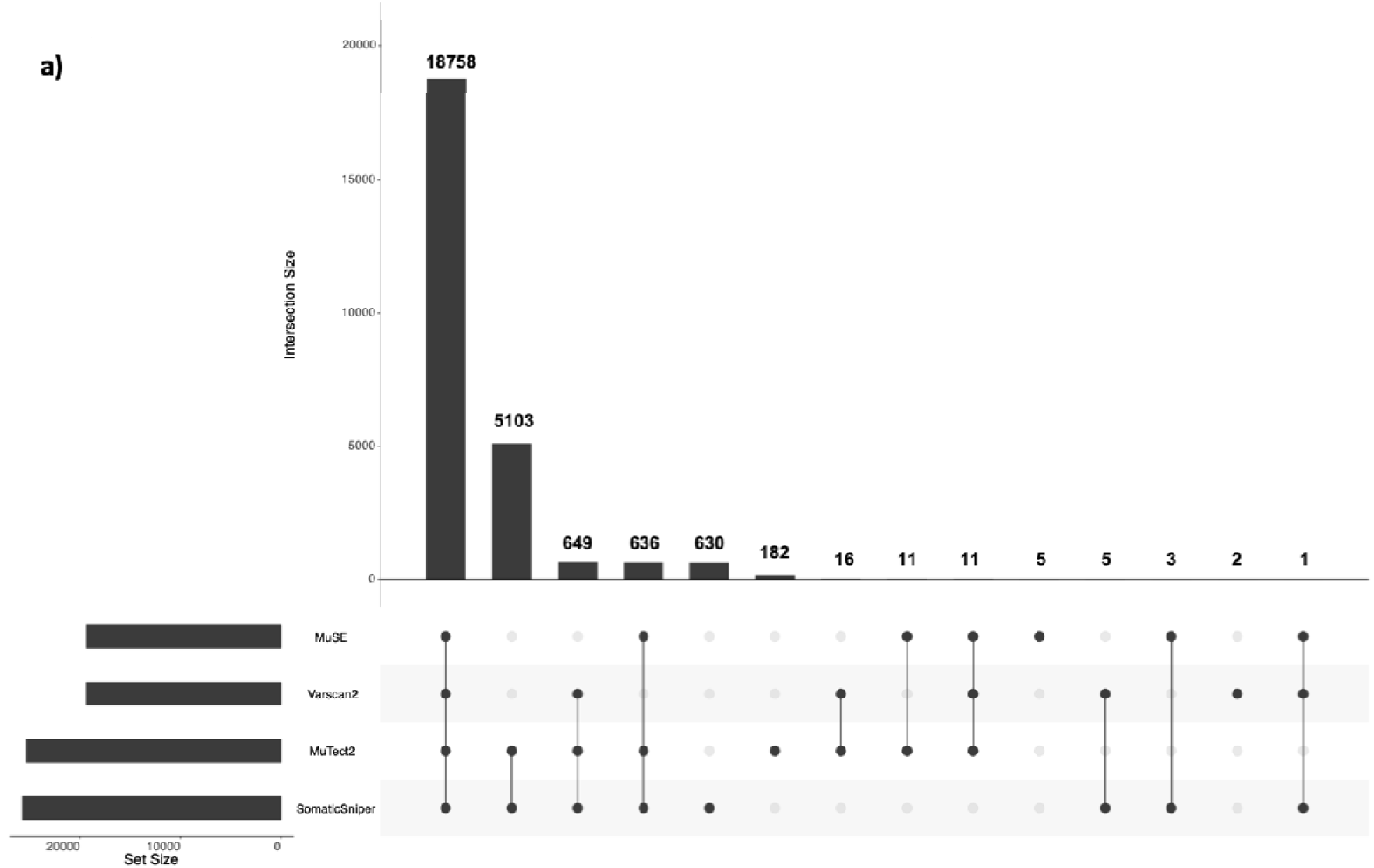
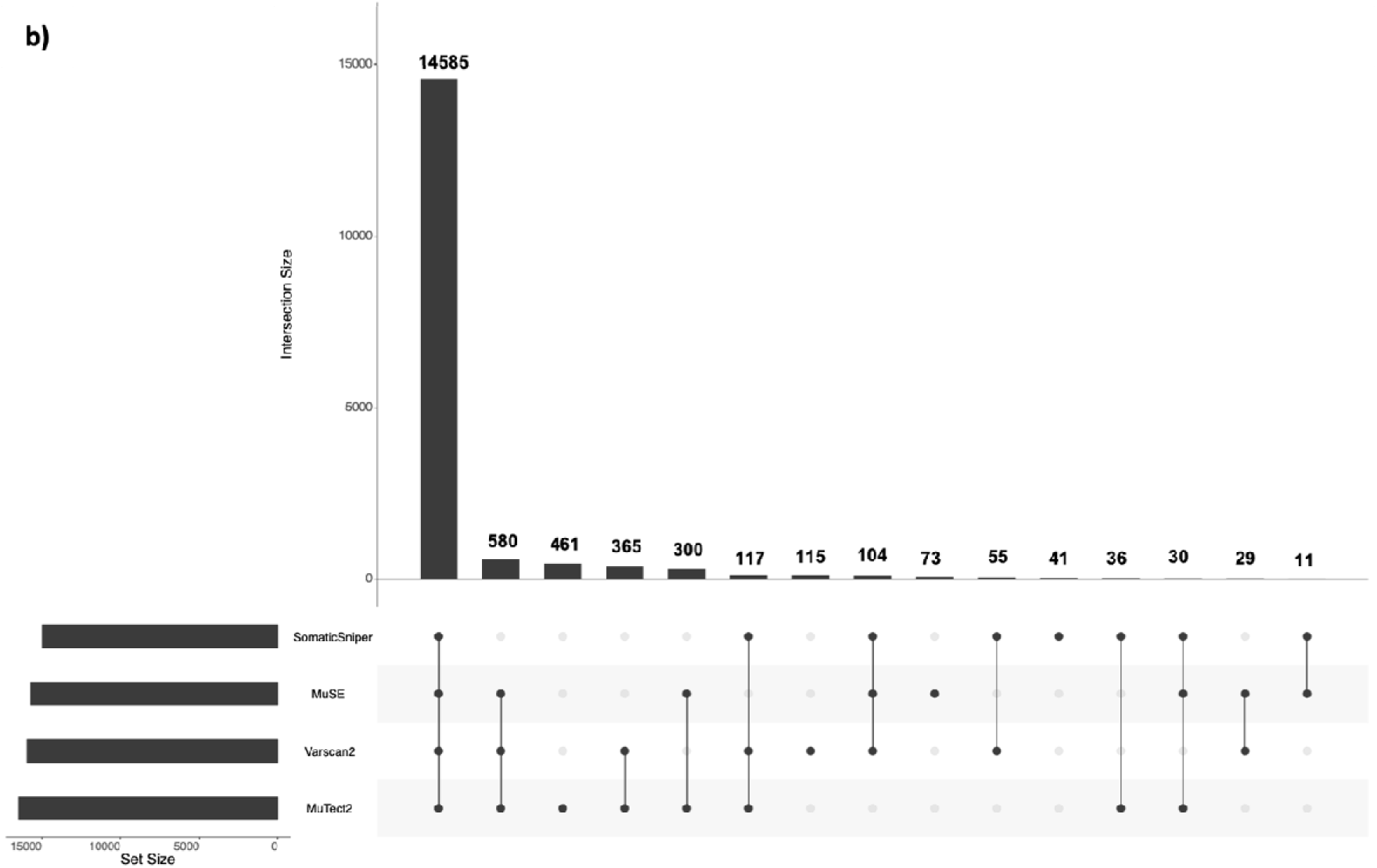
Upset-plot for distribution of genes in four techniques. a) From VCF files b) From MAF files.

### Comparison of Different MAF files

To compare different mutation calling techniques, we have taken processed and annotated MAF files from TCGA. We utilized the Maftools package to comprehensively analyse the somatic variants extracted from MuSE, Mutect2, Varscan2, and SomaticSniper mutation calling technique. From the analysis, we observed few changes in the mutation calling techniques for the same cohort of samples. For example, MuSE and SomaticSniper MAF files (Figure 2A, 2B) only report SNPs on the other side Varscan2, and MuTect2 (Figure 2C, 2D) represent SNPs, INS, and DEL under the variant type.

**Figure 2:**
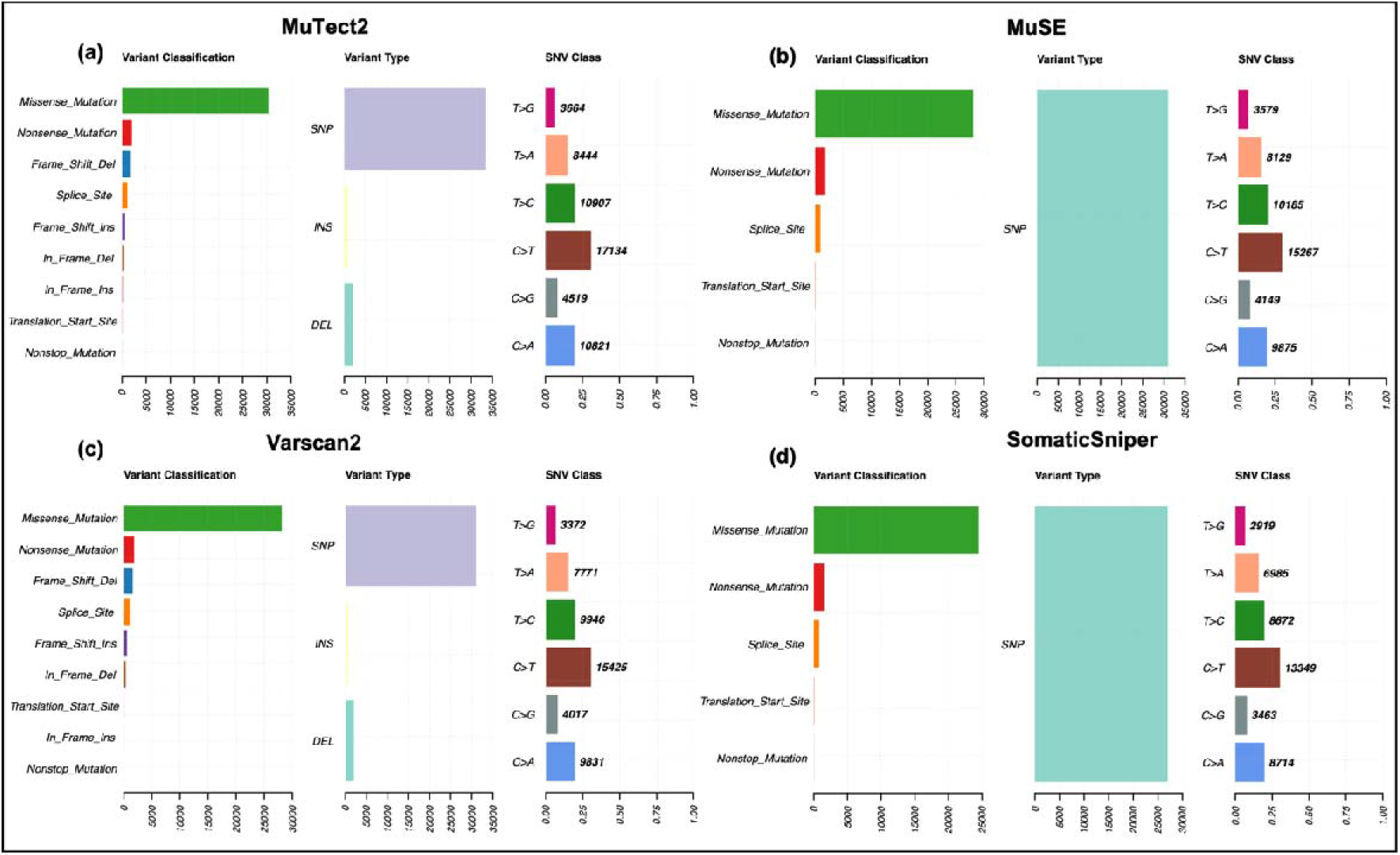
Visualization of mutation summary (variants classification, type and SNVs) for MuTect2, MuSE, Varscan2 and SomaticSniper MAF files.

In Varscan2 and MuTect2, the variant classification distribution represents nine types of mutations such as Missense_Mutation, Nonsense_Mutation, Splice_Site, Translational_Start_Site, Frame_Shift_Ins, Frame_Shift_Del, In_Frame_Ins, In_Frame_Del, and Nonstop_Mutations, while MuSE and SomaticSniper MAF files consist Missense_Mutation, Nonsense_Mutation, Splice_Site, Translational_Start_Site, Nonstop_Mutations. The SNV class visualizes the single-nucleotide variants in the TCGA cohort, we observed that all the methods present diverse distribution of SNV as shown in (Figure 2). Oncoplots generated by the Maftools visualization module illustrating the somatic landscape of the cancer patients for Varscan2, MuTect2, MuSE and SomaticSniper MAF files. In Figure 3, we display the topmost mutated genes with their mutation percentage (>=5%) in total number of samples. From the results we observed that, TP53 is highly mutated gene and have almost 20% or >20% mutations among different techniques.

**Figure 3:**
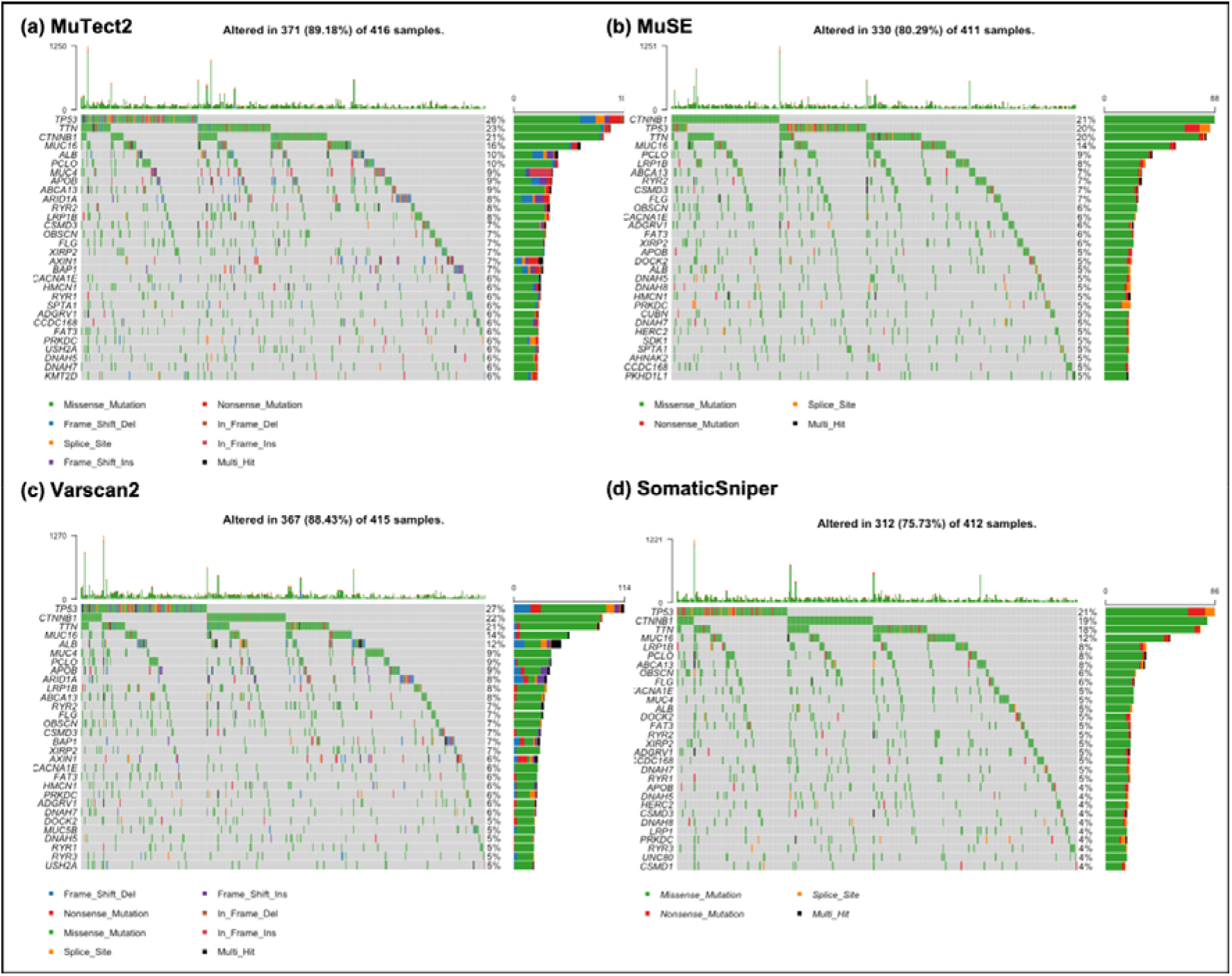
Oncoplot visualization of mutation frequency of top-most mutated genes. The rows represented the genes with % mutations, and columns display the samples. (a) Illustrates the oncoplot of MuTect2 technique and indicates that 89.18% of samples having mutated genes (b) Illustrates the oncoplot of MuSE technique and shows that 80.29% of samples having mutated genes (c) Presents the oncoplot of Varscan2 approach and shows that 88.43% of samples having mutated genes (d) Illustrates the oncoplot of SomaticSniper technique and indicates that 75.73% of samples having alerted/mutated genes

### Correlation Analysis

By implementing the correlation test we ranked the genes and choose top-10 genes having significant negative-correlation coefficients. The procedure is repeated for all the four techniques from MAF and VCF files of liver cancer patients, which lead to 80 genes in total. The complete correlation analysis is provided in Supplementary Table S1.

### Prognostic Biomarkers for High-Risk Prediction

#### Single gene

Univariate survival analysis was performed using cox-proportional hazard model. We have calculated the HR and p-value for ten genes from each technique for VCF files. SomaticSniper technique has achieved the maximum HR value in single gene based analysis with HR_CLDN20_ = 7.06 and p-value 6.62E-07, followed by Varscan2 with HR_FAM160A2_ = 6.81 and p-value 4.01E-05, followed by MuTect2 based VCF file with HR_SNHG10_ = 5.49 and p-value 3.94E-06, and Muse technique has achieved the HR_CLMP_ of 3.01 with p-value 1.67E-05 as shown in Table 2.

**Table 2:** Hazards ratio for top-10 genes from VCF files derived using MuTect2, MuSE, Varscan2, and SomaticSniper technique.

| MuTect2 |  |  |  |  | MuSE |  |  |  |  |
| --- | --- | --- | --- | --- | --- | --- | --- | --- | --- |
| Gene | HR | P-value | 95% CI | C-index | Gene | HR | P-value | 95% CI | C-index |
| SNHG10 | 5.49 | 3.94E-06 | 2.66 - 11.31 | 0.53 | CLMP | 3.01 | 1.67E-05 | 1.82 - 4.97 | 0.54 |
| WIZ | 2.69 | 9.71E-07 | 1.81 - 4.00 | 0.56 | BIRC6 | 2.80 | 4.46E-04 | 1.58 - 4.99 | 0.54 |
| MGAT4EP | 2.49 | 4.46E-04 | 1.50 - 4.15 | 0.54 | LINC02210-CRHR1 | 2.03 | 6.42E-03 | 1.22 - 3.39 | 0.53 |
| LINC00304 | 2.39 | 7.40E-05 | 1.55 - 3.67 | 0.55 | DHX8 | 2.00 | 2.90E-02 | 1.07 - 3.74 | 0.52 |
| CACNG7 | 1.93 | 5.72E-04 | 1.33 - 2.81 | 0.56 | LINC00972 | 1.91 | 9.31E-03 | 1.17 - 3.10 | 0.54 |
| OR52B6 | 1.83 | 1.12E-03 | 1.27 - 2.63 | 0.56 | PAX7 | 1.90 | 8.29E-04 | 1.30 - 2.76 | 0.56 |
| TYK2 | 1.80 | 2.21E-03 | 1.24 - 2.63 | 0.56 | TAS1R2 | 1.61 | 2.63E-02 | 1.06 - 2.44 | 0.53 |
| PIGO | 1.79 | 1.66E-02 | 1.11 - 2.88 | 0.52 | SNTG1 | 1.53 | 3.37E-02 | 1.03 - 2.27 | 0.54 |
| S100A12 | 1.71 | 1.10E-02 | 1.13 - 2.59 | 0.54 | CNTN5 | 1.34 | 2.25E-01 | 0.83 - 2.16 | 0.51 |
| DNAJC9-AS1 | 1.08 | 6.51E-01 | 0.77 - 1.51 | 0.52 | ZNF521 | 1.26 | 2.63E-01 | 0.84 - 1.91 | 0.52 |
| Varscan2 |  |  |  |  | SomaticSniper |  |  |  |  |
| Gene | HR | P-value | 95% CI | C-index | Gene | HR | P-value | 95% CI | C-index |
| FAM160A2 | 6.81 | 4.01E-05 | 2.73 - 17.02 | 0.52 | CLDN20 | 7.06 | 6.62E-07 | 3.27 - 15.2 | 0.53 |
| LOC100420587 | 5.45 | 1.31E-07 | 2.90 - 10.22 | 0.54 | NR2C2AP | 5.17 | 3.16E-05 | 2.38 - 11.2 | 0.52 |
| SPDYA | 3.08 | 7.70E-04 | 1.60 - 5.94 | 0.53 | ATG9B | 3.34 | 2.59E-04 | 1.75 - 6.37 | 0.53 |
| BRSK2 | 2.55 | 1.01E-03 | 1.46 - 4.46 | 0.54 | HAUS5 | 2.79 | 2.22E-05 | 1.74 - 4.48 | 0.55 |
| ADGRF4 | 2.21 | 1.23E-02 | 1.19 - 4.10 | 0.53 | LOC100287329 | 2.58 | 8.23E-04 | 1.48 - 4.49 | 0.53 |
| LINC00972 | 2.11 | 2.18E-03 | 1.31 - 3.41 | 0.55 | P4HTM | 2.18 | 2.43E-02 | 1.11 - 4.31 | 0.52 |
| TM4SF18 | 2.07 | 1.40E-02 | 1.16 - 3.70 | 0.53 | OR6C76 | 2.12 | 1.18E-03 | 1.35 - 3.35 | 0.54 |
| OR5AS1 | 1.86 | 1.43E-02 | 1.13 - 3.06 | 0.54 | CLK2 | 1.94 | 3.58E-02 | 1.05 - 3.61 | 0.52 |
| PDE11A | 1.72 | 2.74E-03 | 1.21 - 2.46 | 0.55 | FAM187B | 1.64 | 1.51E-02 | 1.10 - 2.43 | 0.55 |
| LOC101929073 | 1.29 | 2.98E-01 | 0.80 - 2.11 | 0.52 | NOMO3 | 1.34 | 1.45E-01 | 0.90 - 1.98 | 0.52 |

HR: Hazard ratio; 95% CI: 95% Confidence Interval; C-index: Concordance index

Similar analysis was done for MAF files from each technique and HR values were calculated. As exhibited in Table 3, Mutect2 technique based MAF file has achieved the maximum HR_LAMC3_ = 9.25 with p-value 1.78E-06, followed by Varscan2 with HR_SYDE1_ 8.46 and 3.71E-05, followed by MuSE technique with HR_ITGB8_ 8.30 and p-value 5.69E-07, then followed by SomaticSniper with HR_CAD_ 5.56 and p-value 8.10E-04.

**Table 3:** Hazards ratio for top-10 genes from MAF files derived using MuTect2, MuSE, Varscan2, and SomaticSniper technique.

| MuTect2 |  |  |  |  | MuSE |  |  |  |  |
| --- | --- | --- | --- | --- | --- | --- | --- | --- | --- |
| Gene | HR | P-value | 95% CI | C-index | Gene | HR | P-value | 95% CI | C-index |
| LAMC3 | 9.25 | 1.78E-06 | 3.71 - 23.05 | 0.52 | ITGB8 | 8.37 | 5.69E-07 | 3.64 - 19.24 | 0.52 |
| EVC2 | 4.30 | 8.66E-05 | 2.08 - 8.91 | 0.53 | TBX3 | 8.10 | 6.06E-05 | 2.91 - 22.53 | 0.52 |
| NYNRIN | 3.94 | 1.22E-03 | 1.72 - 9.05 | 0.52 | SIPA1L3 | 4.90 | 5.54E-05 | 2.26 - 10.61 | 0.52 |
| KIAA2026 | 3.85 | 1.49E-03 | 1.68 - 8.86 | 0.52 | CAD | 4.45 | 3.58E-03 | 1.63 - 12.14 | 0.52 |
| SUPT20H | 3.41 | 7.53E-03 | 1.39 - 8.40 | 0.51 | EVC2 | 4.16 | 2.97E-04 | 1.92 - 9.01 | 0.52 |
| BRINP2 | 2.83 | 2.43E-02 | 1.14 - 6.98 | 0.52 | ARHGEF11 | 3.17 | 2.37E-02 | 1.17 - 8.64 | 0.51 |
| LRP1B | 1.93 | 7.81E-03 | 1.19 - 3.14 | 0.54 | BRINP2 | 2.80 | 2.56E-02 | 1.13 - 6.92 | 0.52 |
| TP53 | 1.48 | 3.60E-02 | 1.03 - 2.14 | 0.55 | PCDH15 | 1.72 | 1.20E-01 | 0.87 - 3.39 | 0.51 |
| TG | 1.46 | 4.53E-01 | 0.54 - 3.97 | 0.51 | TG | 1.46 | 4.55E-01 | 0.54 - 3.97 | 0.51 |
| PCDH15 | 1.43 | 3.30E-01 | 0.70 - 2.93 | 0.51 | CSMD3 | 1.24 | 4.54E-01 | 0.71 - 2.15 | 0.51 |
| Varscan2 |  |  |  |  | SomaticSniper |  |  |  |  |
| Gene | HR | P-value | 95% CI | C-index | Gene | HR | P-value | 95% CI | C-index |
| SYDE1 | 8.46 | 3.71E-05 | 3.07 - 23.35 | 0.52 | CAD | 5.56 | 8.10E-04 | 2.04 - 15.17 | 0.52 |
| ALPP | 4.33 | 1.44E-03 | 1.76 - 10.66 | 0.52 | TOP2A | 4.63 | 2.73E-03 | 1.70 - 12.62 | 0.52 |
| KIAA2026 | 3.85 | 1.49E-03 | 1.68 - 8.86 | 0.52 | KIAA2026 | 4.01 | 2.62E-03 | 1.62 - 9.93 | 0.52 |
| CAD | 3.32 | 1.91E-02 | 1.22 - 9.04 | 0.51 | EVC2 | 4.00 | 1.04E-03 | 1.75 - 9.17 | 0.52 |
| BRINP2 | 2.83 | 2.43E-02 | 1.14 - 6.98 | 0.52 | KTN1 | 2.56 | 1.09E-01 | 0.81 - 8.10 | 0.51 |
| TP53 | 1.60 | 9.85E-03 | 1.12 - 2.30 | 0.56 | EPHA3 | 2.25 | 1.67E-01 | 0.71 - 7.13 | 0.51 |
| PCDH15 | 1.48 | 2.81E-01 | 0.72 - 3.05 | 0.51 | KIF26B | 2.03 | 1.66E-01 | 0.74 - 5.55 | 0.51 |
| TG | 1.46 | 4.53E-01 | 0.54 - 3.97 | 0.51 | PCDH15 | 1.76 | 1.78E-01 | 0.77 - 4.02 | 0.51 |
| PLCB1 | 1.25 | 7.00E-01 | 0.40 - 3.96 | 0.50 | TP53 | 1.63 | 1.20E-02 | 1.11 - 2.38 | 0.55 |
| XIRP2 | 1.11 | 7.55E-01 | 0.58 - 2.12 | 0.51 | TG | 1.18 | 8.17E-01 | 0.29 - 4.79 | 0.50 |

HR: Hazard ratio; 95% CI: 95% Confidence Interval; C-index: Concordance index

#### Multiple Gene

In order to explore the effect of mutations in all the selected genes altogether, we have predicted the survival time to estimate the high-risk group in liver cancer patients. Using the predicted OS time, HR and p-value was computed with cox proportional hazard models for each technique corresponds to each file type. We achieved highest HR 4.50 with highly significant p-value 3.83E-15 for the VCF files generated using the MuTect2 technique (Figure 4A). However, in case of MAF files, MuSE technique performed best among other techniques with HR 2.47 and p-value 9.64E-07 (Figure 4B). Additionally, KM survival plots clearly represents the segregation of high- and low-risk groups; the comparison of different mutation calling techniques based on two file formats is shown in Figure 4.

**Figure 4:**
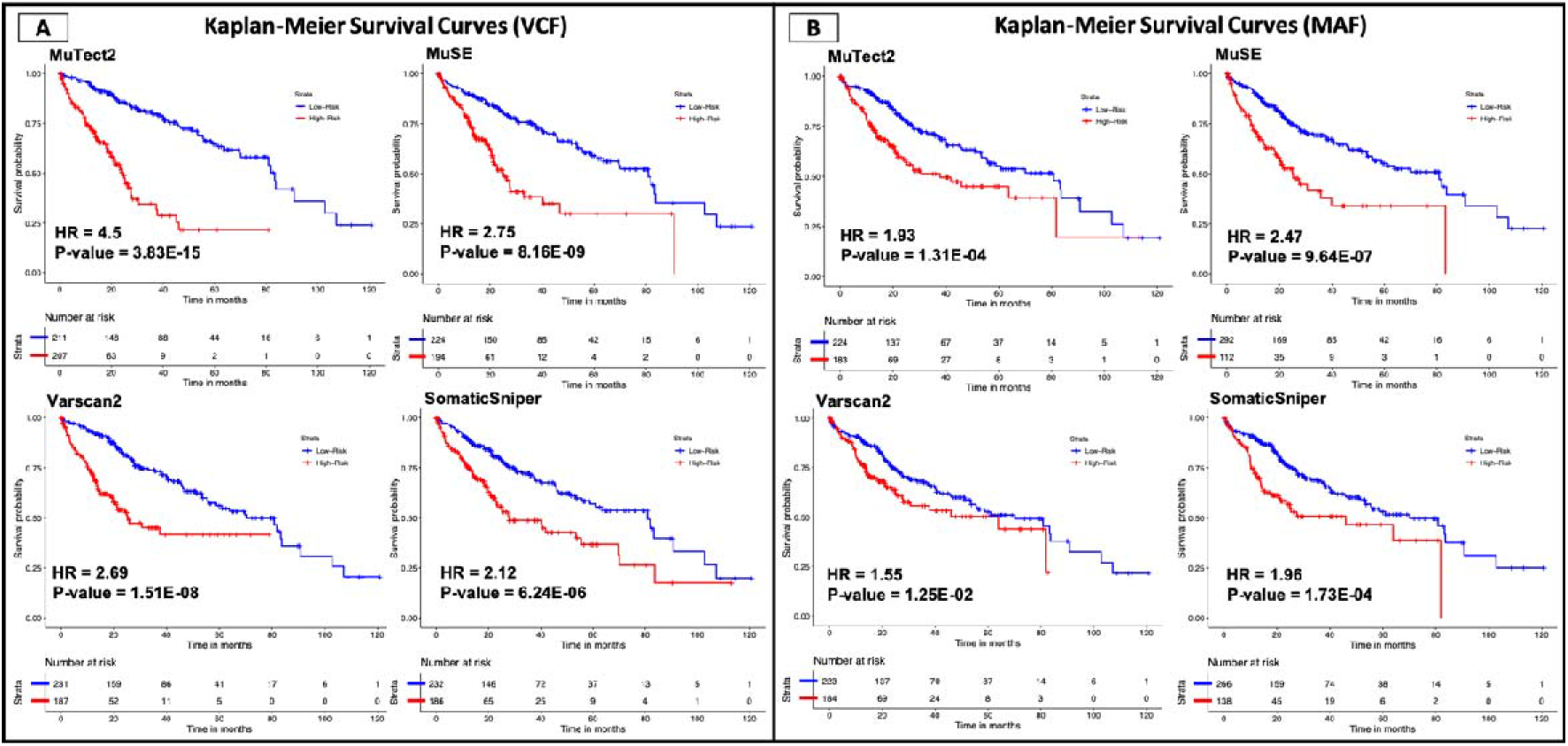
Kaplan Meier survival curves for the risk estimation of liver cancer patients based on the combined effect of mutation (A) survival plots for the VCF files (B) survival plots for the MAF files.

#### Prediction of Overall Survival of Patients

To predict the overall survival for liver cancer patients, we have used number of mutations in the top-10 genes as the input feature and developed regression models for VCF and MAF files for each technique, using seven different regressors such as, Linear (LR), Lasso (LAS), Ridge (RID), Elastic Net (ENT), Decision Tree (DTR), Random Forest (RFR), and Support Vector (SVR). Table 4 exhibits the performance of best performing regressor in each file type. Performance of all the regressors for each file type and technique is reported in Supplementary Table S2. In case of MuTect2 technique, the OS predicted using VCF files have MAE 12.52 and significant correlation of 0.57 between the true and predicted OS; whereas in MAF file the MAE is 16.47 with R 0.37. Whereas, MuSE technique has achieved the minimum MAE of 13.88 and 16.89 along with R of 0.51 and 0.34, for VCF and MAF file respectively. In files generated using Varscan2 technique, for VCF file the minimum MAE is 14.57 with R 0.48, whereas for MAF file it is 16.53 with R 0.36. VCF and MAF file generated using SomaticSniper technique reported minimum MAE of 15.76 (R=0.40) and 16.72 (R=0.33), respectively. As shown in Table 4, for VCF as well as MAF files, MuTect2 technique outperformed the other techniques in terms of MAE, RMSE and R-value.

**Table 4:** Performance of best regressors on top-10 genes from VCF and MAF files extracted using all techniques.

| Technique | File Type | MAE | RMSE | R | p-value |
| --- | --- | --- | --- | --- | --- |
| MuTect2 | VCF | 12.52 | 19.58 | 0.57 | 7.00E-37 |
|  | MAF | 16.47 | 22.16 | 0.37 | 1.31E-14 |
| MuSE | VCF | 13.88 | 20.38 | 0.51 | 1.38E-29 |
|  | MAF | 16.89 | 22.48 | 0.34 | 1.68E-12 |
| Varscan2 | VCF | 14.57 | 20.78 | 0.48 | 4.77E-26 |
|  | MAF | 16.53 | 22.26 | 0.36 | 9.11E-14 |
| SomaticSniper | VCF | 15.76 | 21.82 | 0.40 | 3.31E-17 |
|  | MAF | 16.72 | 22.26 | 0.33 | 8.46E-12 |

MAE: Mean Absolute Error; RMSE: Root Mean Square Error; HR: Hazard Ratio; R: Correlation Coefficient

#### Discrimination of Low- and High-Risk patients

Initially, the dataset was divided into two groups, i.e., the high-risk and low-risk group based on the median OS. Samples with OS time less than the median OS time were designated to the high-risk group, whereas the remaining were assigned to the low-risk group. To assess the ability of the number of mutations/gene/samples to classify the patients into the high and low-risk groups, classification models were developed on top 10 genes for each technique and file type, using eight different classifiers such as RF, LR, XGB, DT, KNN, GNB, ET and SVC. The performance of all the classifiers for every model generated on each technique for both the files are reported in Supplementary Table S3.

Number of mutations reported through each technique were used to develop models to predict the high- and low-risk group. In case of VCF file derived using Mutect2, SVC-based model achieved AUROC of 0.72 and 0.69 in training and validation data, respectively as shown in Table 5. Similarly, ET-based model developed on genes from MAF files extracted using MuTect2 technique performed with AUROC of 0.57 and 0.67 on training and validation dataset, respectively. For MuSE technique, GNB-based model developed on genes from VCF files achieved AUROC of 0.66 and 0.68 on training and validation data whereas, ET-based model developed on genes from MAF files achieved 0.60 and 0.51 AUROC on training and validation dataset, respectively. For the genes obtained from the Varscan2 technique, SVC-based model with genes from VCF file performed best with AUROC 0.68 and 0.64 on the training and validation dataset, with the minimum difference in sensitivity and specificity, whereas for MAF files, LR-based model achieved AUROC of 0.63 and 0.63 on training and validation dataset. For SomaticSniper technique, LR-based model developed on genes from VCF files achieved AUROC of 0.63 and 0.65 on training and validation data whereas, LR-based model developed on genes from MAF files achieved 0.60 and 0.64 AUROC on training and validation dataset, respectively. For VCF as well as MAF files, MuTect2 technique performed best among other techniques in terms of difference between sensitivity and specificity as well as AUROC.

**Table 5:**
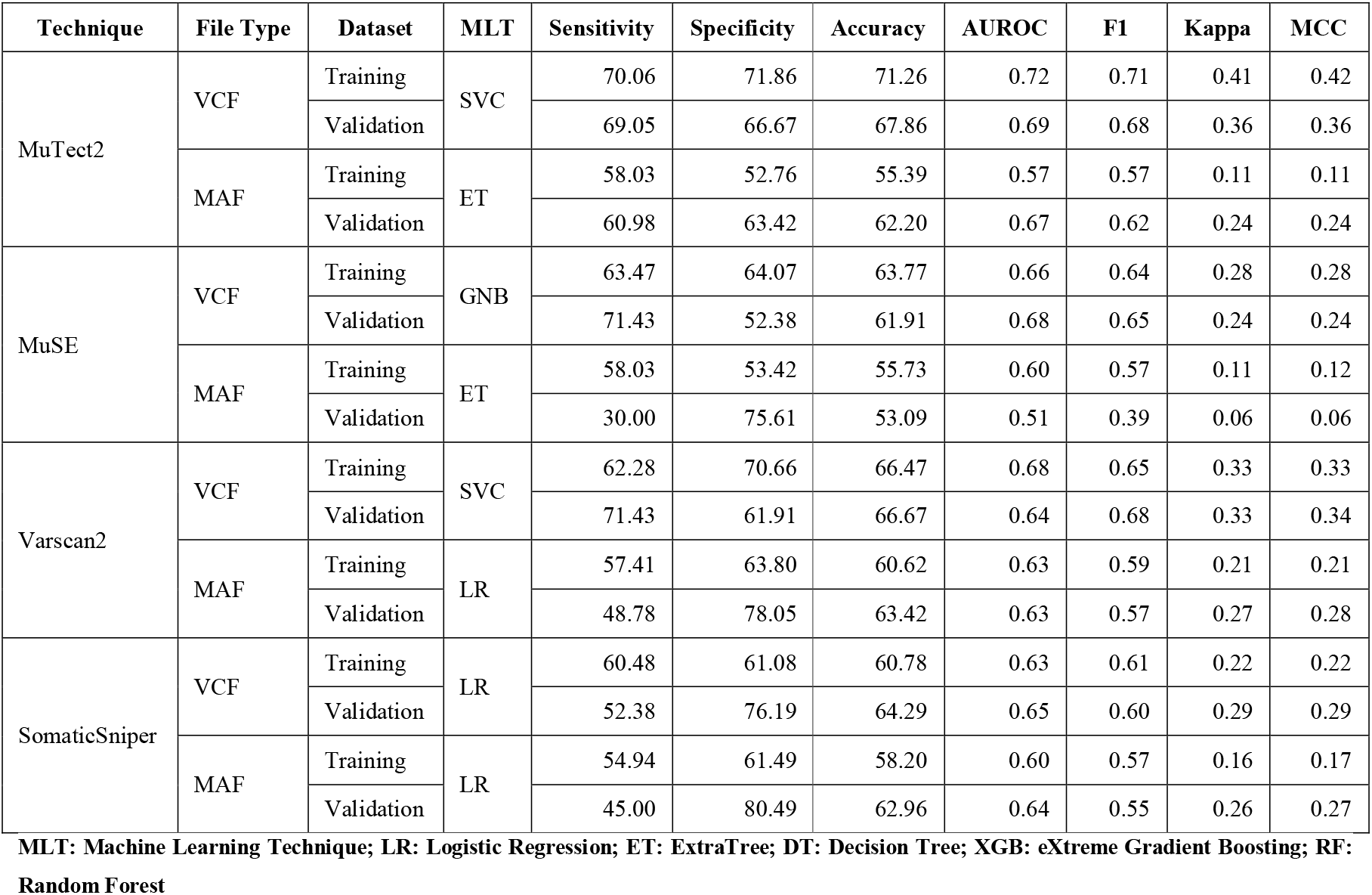
Performance of best classifiers on top-10 genes from VCF and MAF files extracted using all techniques.

| Technique | File Type | Dataset | MLT | Sensitivity | Specificity | Accuracy | AUROC | F1 | Kappa | MCC |
| --- | --- | --- | --- | --- | --- | --- | --- | --- | --- | --- |
| MuTect2 | VCF | Training | SVC | 70.06 | 71.86 | 71.26 | 0.72 | 0.71 | 0.41 | 0.42 |
|  |  | Validation |  | 69.05 | 66.67 | 67.86 | 0.69 | 0.68 | 0.36 | 0.36 |
|  | MAF | Training | ET | 58.03 | 52.76 | 55.39 | 0.57 | 0.57 | 0.11 | 0.11 |
|  |  | Validation |  | 60.98 | 63.42 | 62.20 | 0.67 | 0.62 | 0.24 | 0.24 |
| MuSE | VCF | Training | GNB | 63.47 | 64.07 | 63.77 | 0.66 | 0.64 | 0.28 | 0.28 |
|  |  | Validation |  | 71.43 | 52.38 | 61.91 | 0.68 | 0.65 | 0.24 | 0.24 |
|  | MAF | Training | ET | 58.03 | 53.42 | 55.73 | 0.60 | 0.57 | 0.11 | 0.12 |
|  |  | Validation |  | 30.00 | 75.61 | 53.09 | 0.51 | 0.39 | 0.06 | 0.06 |
| Varscan2 | VCF | Training | SVC | 62.28 | 70.66 | 66.47 | 0.68 | 0.65 | 0.33 | 0.33 |
|  |  | Validation |  | 71.43 | 61.91 | 66.67 | 0.64 | 0.68 | 0.33 | 0.34 |
|  | MAF | Training | LR | 57.41 | 63.80 | 60.62 | 0.63 | 0.59 | 0.21 | 0.21 |
|  |  | Validation |  | 48.78 | 78.05 | 63.42 | 0.63 | 0.57 | 0.27 | 0.28 |
| SomaticSniper | VCF | Training | LR | 60.48 | 61.08 | 60.78 | 0.63 | 0.61 | 0.22 | 0.22 |
|  |  | Validation |  | 52.38 | 76.19 | 64.29 | 0.65 | 0.60 | 0.29 | 0.29 |
|  | MAF | Training | LR | 54.94 | 61.49 | 58.20 | 0.60 | 0.57 | 0.16 | 0.17 |
|  |  | Validation |  | 45.00 | 80.49 | 62.96 | 0.64 | 0.55 | 0.26 | 0.27 |

MLT: Machine Learning Technique; LR: Logistic Regression; ET: ExtraTree; DT: Decision Tree; XGB: eXtreme Gradient Boosting; RF: Random Forest

## Discussion

Liver cancer is a global problem and occurs after severe liver diseases [31]. Chronic liver diseases are associated with cancer development and prompt progressive mutations at the genomic level [32, 33]. Previous studies report that liver cancer is associated with poor prognosis and a high mortality rate amongst the most frequent cancer types [34, 35]. Nowadays, several mutation calling techniques are available to identify the mutation landscape in tumor/normal patients. Hitherto, there is not an appropriate comparison of mutation detection methods for the predictive and prognostic analysis. In this study, we examine the performance of four widely used mutation calling techniques such as MuTect2, MuSE, Varscan2, and SomaticSniper using TCGA liver cancer cohort. We have applied various techniques in order to compare all the methods for predicting and analysing prognostic biomarkers in liver cancer patients. First, we have used VCF and MAF files generated by the different mutation calling methods. We have used the most popular methods (ANNOVAR and Maftools) to identify the gene-associated mutations in liver cancer samples. Further, we observed that the VCF files of Mutect2 and SomaticSniper report highest number of mutated genes and cover over 5 million mutations. Whereas, MAF files reports comparatively less mutated genes for each technique as shown in Table 1.

Then, we performed correlation analysis in order to understand the impact of mutations on the survival of liver cancer patients. On performing the univariate survival analysis on VCF files, we observed that LncRNA SNGH10, CLMP, FAM160A2 and CLDN20 achieved the highest HR value in MuTect2, MuSE, Varscan2 and SomaticSniper technique respectively. As shown by Lan et al. LncRNA SNGH10 is an oncogenic lncRNA in liver cancer patients and reduces the survival of the patients [36]. It’s down-regulation is also associated with the poor survival non-small cell lung cancer with HR 2.09 with p-value 0.02 [37]. Our study also corresponds with the previous studies and exhibits that the mutations in SNGH10 gene is associated with poor outcome in liver cancer patients with HR 5.49 and p-value 3.94E-06. Whereas, the differential expression of CLMP gene is associated with the progression of cancers of the breast cancer [38]. Yang et al. also reported the significance of CLDN20 gene in the survival of breast cancer patients with HR 1.38 and p-value 0.047 [39]. However, our analysis reveal the role of CLMP and CLDN20 gene in the survival of liver cancer patients. Further, in case of MAF files, the univariate survival analysis reveals that SYDE1, LAMC3, ITGB8, CAD, EVC2, NYNRIN, BRSK2, TP53 genes significantly reduces the overall survival. As shown by the recent study that SYDE1 act as an oncogene and overexpressed in glioma patients makes it an important diagnostic and prognostic biomarker [40]. Moreover, the down-regulation of LAMC3 is correlated with the poor prognosis and metastasis in the ovarian cancer patients [41]. A study also reveals that mutations associated with LAMC3 genes may cause PNH (a rare disorder of clonal stem cell in foetus), which may leads high mortality rate infection and premature birth [42, 43]. We also observed that mutations associated with LAMC3 significantly reduces the survival of patients with HR = 9.25 and p-value 1.78E-06. In addition, ITGB8 is shown to be highly upregulated in high-grade ovarian cancer patients, which leads to shorter OS with significant HR 1.42 [44]. Paul et.al, also reveals that EVC2 gene is highly mutated in breast cancer patients and dysregulates pathways like (mTOR, CDK/RB, cAMP/PKA, WNT, etc) [45]. Our study show that mutations associated with EVC2 genes reduces the overall survival of patients with HR = 4.3 and p-value 8.66E-05. Researchers have shown that the overexpression of BRSK2 gene correlated with the patients survival and prognosis in pancreatic cancer [46]. Of Note, several studies reports that TP53 is the highly mutated gene among most of the human cancers and affects the survival of cancer patients [47–51]. In current study, we also found that TP53 is the highly mutated gene among the liver cancer patients and covers almost 20% mutations. Correlation and survival analysis shown that mutation associated with TP53 significantly reduces the overall survival with HR = 1.63 and p-value 1.20E-02 among liver cancer patients. While considering the combined effect of selected genes in each file, MuTect2 technique outperformed all the other techniques in VCF file with HR 4.50 (p-value 3.83E-15), whereas MuSE technique outperformed other mutation calling methods with HR 2.47 (p-value 9.64E-07) in case of MAF files (Figure 4).

Furthermore, to compare the different mutation calling techniques we develop various survival prediction and classification models using the top-10 genes respective to each file type (Table 4 and 5). The predicted survival time employed for the stratification of high-risk and low-risk groups. Models based on ten selected genes from VCF file of MuTect2 technique performed best among the other techniques in stratification of patients in high- and low-risk group, as well as in OS time prediction. Our findings suggest that the VCF file generated using MuTect2 mutation calling technique provides the comprehensive information which can be used for the risk-estimation of liver cancer cohort. Furthermore, this needs to be confirmed on the other cancer cohorts to explore the prognostic potential of mutations.

## Declarations

### Funding

The current work has not received any specific grant from any funding agencies.

### Conflict of Interests

The authors declare no competing financial and non-financial interests.

### Ethics Approval

Not applicable

### Consent to participate

Not applicable

### Conflict of Publication

Not applicable

## Acknowledgements

Authors are thankful to the Department of Computational Biology, IIIT-Delhi for infrastructure, Department of Biotechnology (DBT), Department of Science and Technology (DST-INSPIRE) for financial support and fellowships.

## Author contribution

SP, AD, and GPSR collected and processed the datasets. SP, AD, and GPSR implemented the algorithms. SP, AD, and GPSR developed the prediction models. SP, AD, and GPSR analyzed the results. SP, AD, and GPSR penned the manuscript. GPSR conceived and coordinated the project and provided overall supervision to the project. All authors have read and approved the final manuscript.

## Notes

### Competing Interest Statement

The authors have declared no competing interest.

## References

1. Sung H, Ferlay J, Siegel RL et al. Global Cancer Statistics 2020: GLOBOCAN Estimates of Incidence and Mortality Worldwide for 36 Cancers in 185 Countries, CA Cancer J Clin 2021;71:209–249.

2. Gerlinger M, Rowan AJ, Horswell S et al. Intratumor heterogeneity and branched evolution revealed by multiregion sequencing, N Engl J Med 2012;366:883–892.

3. Taylor SS. Protein kinases: a diverse family of related proteins, Bioessays 1987;7:24–29.

4. Flaherty KT, Puzanov I, Kim KB et al. Inhibition of mutated, activated BRAF in metastatic melanoma, N Engl J Med 2010;363:809–819.

5. Lynch TJ, Bell DW, Sordella R et al. Activating mutations in the epidermal growth factor receptor underlying responsiveness of non-small-cell lung cancer to gefitinib, N Engl J Med 2004;350:2129–2139.

6. Hirsch FR, Scagliotti GV, Mulshine JL et al. Lung cancer: current therapies and new targeted treatments, Lancet 2017;389:299–311.

7. Audeh MW, Carmichael J, Penson RT et al. Oral poly(ADP-ribose) polymerase inhibitor olaparib in patients with BRCA1 or BRCA2 mutations and recurrent ovarian cancer: a proof-of-concept trial, Lancet 2010;376:245–251.

8. LaDuca H, Farwell KD, Vuong H et al. Exome sequencing covers >98% of mutations identified on targeted next generation sequencing panels, PLoS One 2017;12:e0170843.

9. Lelieveld SH, Spielmann M, Mundlos S et al. Comparison of Exome and Genome Sequencing Technologies for the Complete Capture of Protein-Coding Regions, Hum Mutat 2015;36:815–822.

10. Hartley T, Wagner JD, Warman-Chardon J et al. Whole-exome sequencing is a valuable diagnostic tool for inherited peripheral neuropathies: Outcomes from a cohort of 50 families, Clin Genet 2018;93:301–309.

11. Koboldt DC, Zhang Q, Larson DE et al. VarScan 2: somatic mutation and copy number alteration discovery in cancer by exome sequencing, Genome Res 2012;22:568–576.

12. Kim S, Scheffler K, Halpern AL et al. Strelka2: fast and accurate calling of germline and somatic variants, Nat Methods 2018;15:591–594.

13. Alioto TS, Buchhalter I, Derdak S et al. A comprehensive assessment of somatic mutation detection in cancer using whole-genome sequencing, Nat Commun 2015;6:10001.

14. do Valle IF, Giampieri E, Simonetti G et al. Optimized pipeline of MuTect and GATK tools to improve the detection of somatic single nucleotide polymorphisms in whole-exome sequencing data, BMC Bioinformatics 2016;17:341.

15. Cibulskis K, Lawrence MS, Carter SL et al. Sensitive detection of somatic point mutations in impure and heterogeneous cancer samples, Nat Biotechnol 2013;31:213–219.

16. Fan Y, Xi L, Hughes DS et al. MuSE: accounting for tumor heterogeneity using a sample-specific error model improves sensitivity and specificity in mutation calling from sequencing data, Genome Biol 2016;17:178.

17. Larson DE, Harris CC, Chen K et al. SomaticSniper: identification of somatic point mutations in whole genome sequencing data, Bioinformatics 2012;28:311–317.

18. Revathidevi S, Munirajan AK. Akt in cancer: Mediator and more, Semin Cancer Biol 2019;59:80–91.

19. Wang K, Li M, Hakonarson H. ANNOVAR: functional annotation of genetic variants from high-throughput sequencing data, Nucleic Acids Res 2010;38:e164.

20. Mayakonda A, Lin DC, Assenov Y et al. Maftools: efficient and comprehensive analysis of somatic variants in cancer, Genome Res 2018;28:1747–1756.

21. Grossman RL, Heath AP, Ferretti V et al. Toward a Shared Vision for Cancer Genomic Data, N Engl J Med 2016;375:1109–1112.

22. Wei L, Jin Z, Yang S et al. TCGA-assembler 2: software pipeline for retrieval and processing of TCGA/CPTAC data, Bioinformatics 2018;34:1615–1617.

23. Goel MK, Khanna P, Kishore J. Understanding survival analysis: Kaplan-Meier estimate, Int J Ayurveda Res 2010;1:274–278.

24. Pedregosa F, Varoquaux G, Gramfort A et al. Scikit-learn: Machine Learning in Python, Journal of Machine Learning Research 2012;12:2825–2830.

25. Patiyal S, Agrawal P, Kumar V et al. NAGbinder: An approach for identifying N-acetylglucosamine interacting residues of a protein from its primary sequence, Protein Sci 2020;29:201–210.

26. Kaur H, Dhall A, Kumar R et al. Identification of Platform-Independent Diagnostic Biomarker Panel for Hepatocellular Carcinoma Using Large-Scale Transcriptomics Data, Front Genet 2019;10:1306.

27. Dhall A, Patiyal S, Kaur H et al. Computing Skin Cutaneous Melanoma Outcome From the HLA-Alleles and Clinical Characteristics, Front Genet 2020; 11:221.

28. Bhalla S, Kaur H, Dhall A et al. Prediction and Analysis of Skin Cancer Progression using Genomics Profiles of Patients, Sci Rep 2019;9:15790.

29. Schemper M. The relative importance of prognostic factors in studies of survival, Stat Med 1993;12:2377–2382.

30. Lex A, Gehlenborg N, Strobelt H et al. UpSet: Visualization of Intersecting Sets, IEEE Trans Vis Comput Graph 2014;20:1983–1992.

31. Davis GL, Dempster J, Meler JD et al. Hepatocellular carcinoma: management of an increasingly common problem, Proc (Bayl Univ Med Cent) 2008;21:266–280.

32. Muller M, Bird TG, Nault JC. The landscape of gene mutations in cirrhosis and hepatocellular carcinoma, J Hepatol 2020;72:990–1002.

33. Farazi PA, DePinho RA. Hepatocellular carcinoma pathogenesis: from genes to environment, Nat Rev Cancer 2006;6:674–687.

34. Lin L, Yan L, Liu Y et al. The Burden and Trends of Primary Liver Cancer Caused by Specific Etiologies from 1990 to 2017 at the Global, Regional, National, Age, and Sex Level Results from the Global Burden of Disease Study 2017, Liver Cancer 2020;9:563–582.

35. Balogh J, Victor D, 3rd, Asham EH et al. Hepatocellular carcinoma: a review, J Hepatocell Carcinoma 2016;3:41–53.

36. Lan T, Yuan K, Yan X et al. LncRNA SNHG10 Facilitates Hepatocarcinogenesis and Metastasis by Modulating Its Homolog SCARNA13 via a Positive Feedback Loop, Cancer Res 2019;79:3220–3234.

37. Liang M, Wang L, Cao C et al. LncRNA SNHG10 is downregulated in non-small cell lung cancer and predicts poor survival, BMC Pulm Med 2020;20:273.

38. Nilchian A, Johansson J, Ghalali A et al. CXADR-Mediated Formation of an AKT Inhibitory Signalosome at Tight Junctions Controls Epithelial-Mesenchymal Plasticity in Breast Cancer, Cancer Res 2019;79:47–60.

39. Yang G, Jian L, Chen Q. Comprehensive analysis of expression and prognostic value of the claudin family in human breast cancer, Aging (Albany NY) 2021;13:8777–8796.

40. Han Z, Zhuang X, Yang B et al. SYDE1 Acts as an Oncogene in Glioma and has Diagnostic and Prognostic Values, Front Mol Biosci 2021;8:714203.

41. Lei SM, Liu X, Xia LP et al. [Relationships between decreased LAMC3 and poor prognosis in ovarian cancer], Zhonghua Fu Chan Ke Za Zhi 2021;56:489–497.

42. De Angelis C, Byrne AB, Morrow R et al. Compound heterozygous variants in LAMC3 in association with posterior periventricular nodular heterotopia, BMC Med Genomics 2021;14:64.

43. Qian X, Liu X, Zhu Z et al. Variants in LAMC3 Causes Occipital Cortical Malformation, Front Genet 2021;12:616761.

44. He J, Liu Y, Zhang L et al. Integrin Subunit beta 8 (ITGB8) Upregulation Is an Independent Predictor of Unfavorable Survival of High-Grade Serous Ovarian Carcinoma Patients, Med Sci Monit 2018;24:8933–8940.

45. Paul MR, Pan TC, Pant DK et al. Genomic landscape of metastatic breast cancer identifies preferentially dysregulated pathways and targets, J Clin Invest 2020;130:4252–4265.

46. W. Lou Dr. GN. BRSK2 expression as a prognosis marker in pancreatic cancer patients, Journal of Clinical Oncology 2009.

47. Olivier M, Hollstein M, Hainaut P. TP53 mutations in human cancers: origins, consequences, and clinical use, Cold Spring Harb Perspect Biol 2010;2:a001008.

48. Petitjean A, Achatz MI, Borresen-Dale AL et al. TP53 mutations in human cancers: functional selection and impact on cancer prognosis and outcomes, Oncogene 2007;26:2157–2165.

49. Monti P, Menichini P, Speciale A et al. Heterogeneity of TP53 Mutations and P53 Protein Residual Function in Cancer: Does It Matter?, Front Oncol 2020;10:593383.

50. Ungerleider NA, Rao SG, Shahbandi A et al. Breast cancer survival predicted by TP53 mutation status differs markedly depending on treatment, Breast Cancer Res 2018;20:115.

51. Rosenberg S, Okamura R, Kato S et al. Survival Implications of the Relationship between Tissue versus Circulating Tumor DNA TP53 Mutations-A Perspective from a Real-World Precision Medicine Cohort, Mol Cancer Ther 2020;19:2612–2620.

